# Selective inhibition of respiratory complex I reveals a bioenergetic vulnerability in *Francisella*

**DOI:** 10.64898/2026.06.04.730185

**Authors:** Nimra Khalid, Chloe M. Van Horn, Danmeng Li, David A. Ostrov, Aria Eshraghi

## Abstract

*F. tularensis* is a highly infectious Gram-negative bacterial pathogen that causes tularemia, a re-emerging zoonosis of public health concern. Here we identify respiratory complex I as a selective vulnerability in *Francisella* and define the mechanism of action of a pyrazole compound, tolfenpyrad, with species-specific antibacterial activity. Using *F. novicida* as a surrogate model, we demonstrated that tolfenpyrad selectively inhibits growth with no measurable effect on *E. coli* or *P. aeruginosa*. Tolfenpyrad rapidly suppressed oxygen consumption, depleted ATP, collapsed proton motive force, and induced reactive oxygen species, indicating disruption of bacterial metabolism. Biochemical assays demonstrated selective inhibition of NADH-dependent respiration and membrane-associated NADH oxidation, whereas succinate-driven respiration was unaffected. Moreover, the alternative NADH dehydrogenase (*ndh*) was not required for tolfenpyrad activity. Structural docking identified a potential tolfenpyrad-binding pocket within the membrane subunit NuoM. These findings reveal species-specific inhibition of *Francisella* complex I and establish respiratory metabolism as a promising antimicrobial target in these bacteria.

## INTRODUCTION

*Francisella* species are highly infectious Gram-negative bacteria that cause the zoonotic disease tularemia. Of the four recognized *Francisella* subspecies, *F. tularensis* subsp. *tularensis* (Type A) is the most pathogenic and can be transmitted with fewer than 10 organisms, posing a substantial public health risk^1–3^. Transmission to humans occurs through aerosols, arthropod vectors such as ticks and biting flies, direct contact with infected animals, and ingestion of contaminated water or food^4–6^. Since 2010, tularemia has been reported in 47 of the 50 US states with a 57% increase in incidence from the preceding decade^7^. In Europe, tularemia incidence is rising sharply, with Sweden reporting a 20-fold increase in cases over the past decade compared to the preceding ten years, and an 89% increase in cases across the European Union from 2022 to 2023^8–10^. Although antimicrobial therapy is generally effective, delayed treatment increases disease severity, and relapse or treatment failure can occur, particularly in vulnerable populations^11,12^. Untreated infections can result in mortality rates of 30-60%, and there is an ever-present threat of *Francisella* developing resistance to existing antibiotics^6^. These challenges underscore the need for new therapeutic strategies and drug targets for tularemia.

Bacterial metabolism is an attractive target for antimicrobial development due to its central role in energy production and redox homeostasis^13,14^. In many bacteria, ATP is generated through oxidative phosphorylation, in which electrons derived from metabolic substrates are transferred through the electron transport chain (ETC) to generate a proton motive force that drives ATP synthesis^15,16^. Accordingly, small molecules that disrupt respiratory electron transport have the potential to exhibit potent antibacterial activity, as demonstrated in *Mycobacterium tuberculosis*^17,18^. While respiratory metabolism has been characterized extensively in model organisms such as *Escherichia coli*, comparatively little is known about the organization and function of the ETC in *Francisella* or its potential as a therapeutic target^19^. Genomic analyses predict that *Francisella* encodes a branched respiratory chain containing NADH dehydrogenases, quinone carriers, and terminal oxidases, but the functional architecture and vulnerabilities of this system remain poorly defined^20,21^.

Given the importance of the ETC in *Francisella* bioenergetics, we investigated whether small-molecule inhibition of this pathway could impair bacterial growth. We previously identified a pyrazole compound tolfenpyrad as a potent inhibitor of *Francisella* growth; however, its mechanism of action was unclear. Notably, tolfenpyrad is ineffective against other Gram-negative or Gram-positive bacteria, indicating a species-specific vulnerability^22,23^. We recently found that tolfenpyrad antagonizes antibiotics that depend on proton motive force for uptake, suggesting disruption of ETC-linked energetics^24^. Together, these observations support respiratory metabolism as a likely target in *Francisella*.

Here we demonstrate that tolfenpyrad disrupts the ETC in *Francisella*. We find that tolfenpyrad inhibits NADH-dependent electron transport, leading to collapse of proton motive force, ATP depletion, and an increase in reactive oxygen species. Mechanistically, tolfenpyrad targets the NADH dehydrogenase (complex I) activity within the *Francisella* ETC while sparing analogous enzymes in other Gram-negative bacteria. These findings reveal a previously unrecognized vulnerability in *Francisella* bioenergetics and establish respiratory metabolism as a promising therapeutic target.

## RESULTS

### Tolfenpyrad selectively inhibits growth and cellular energetics in Francisella novicida

Building on our previous findings that suggest a genus-specific mechanism that exploits unique physiological features within *Francisella*, we investigated the basis of this selectivity using *Francisella tularensis* subspecies *novicida* (*F. novicida*), a closely related surrogate of *F. tularensis* that can be studied under biosafety level 2 conditions^22,25^. We compared the effects of tolfenpyrad on *F. novicida* to *Escherichia coli* and *Pseudomonas aeruginosa*. All species exhibited comparable growth prior to treatment (Fig. 1A-1C). Upon drug exposure, *E. coli* and *P. aeruginosa* remained unaffected, whereas *F. novicida* underwent pronounced bacteriostasis. These findings are consistent with our prior observations that tolfenpyrad selectively inhibits *Francisella* and indicate that drug sensitivity reflects species-specific physiology rather than generalized toxicity^22^.

**Fig. 1.**
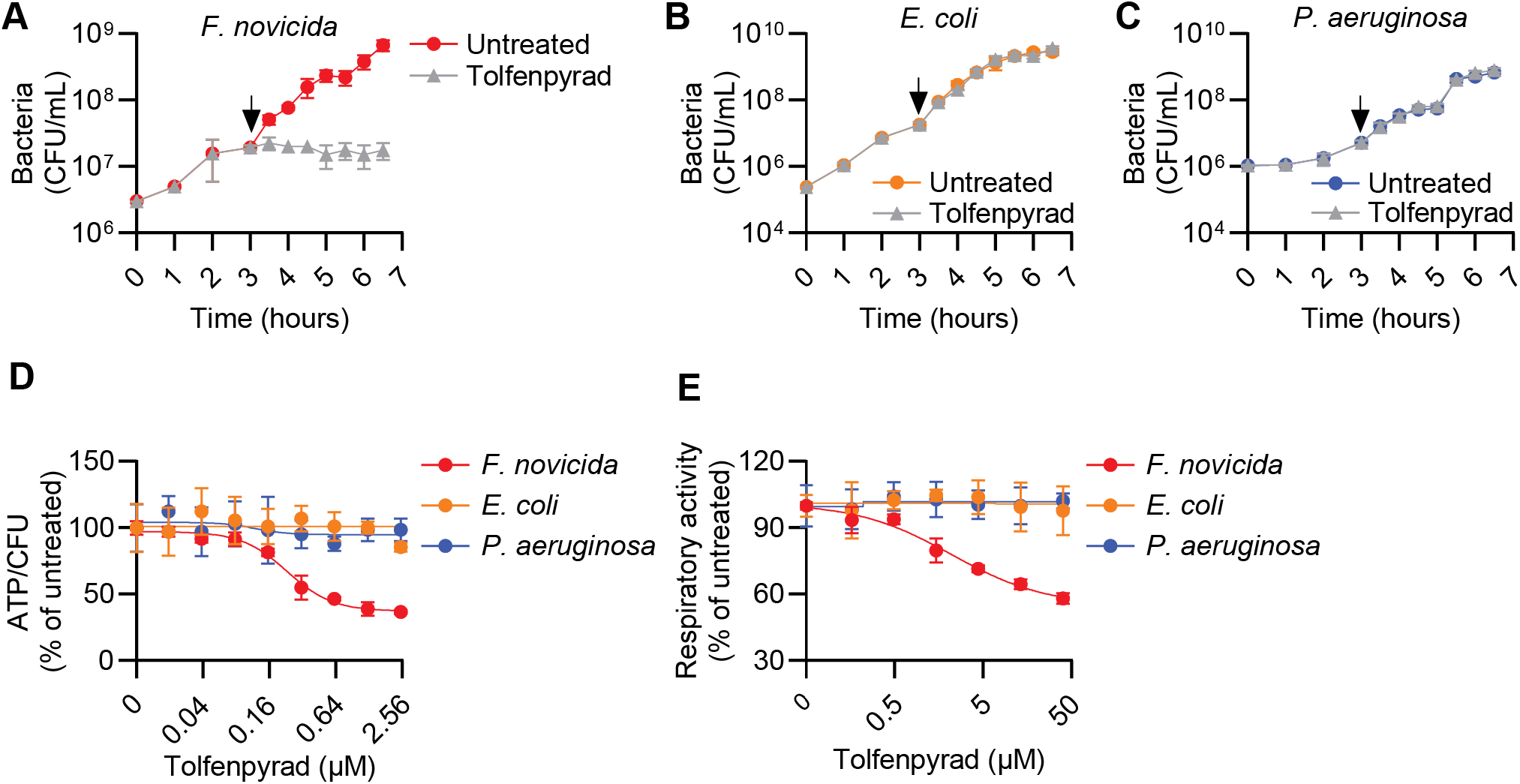
Tolfenpyrad selectively inhibits respiratory activity in *F. novicida*. **(A-C)** Growth kinetics of *F. novicida* **(a)**, *E. coli* **(b)**, and *P. aeruginosa* **(c)** following treatment with 10 µM tolfenpyrad. Arrows indicate the time of tolfenpyrad addition. **(d)** ATP levels normalized to CFU following 45 min treatment with the indicated concentrations of tolfenpyrad. **(e)** Respiratory activity measured by MTT reduction following 45 min treatment with the indicated concentrations of tolfenpyrad. Data are presented as mean ± S.D. and are representative of three or more independent experiments performed with at least three technical replicates per condition.

Given that bacteriostasis often reflects impaired energy metabolism, we quantified intracellular ATP and normalized to viable bacterial counts. Tolfenpyrad selectively reduced intracellular ATP levels in *F. novicida* in a dose-dependent manner and did not measurably alter ATP levels in *E. coli* or *P. aeruginosa* (Fig. 1D). Importantly, normalization to CFU demonstrated that tolfenpyrad reduces ATP production on a per-cell basis rather than through loss of viability. These data suggested that the compound perturbs energy generation rather than bacterial survival directly.

To determine whether ATP depletion reflected impaired respiration, we measured global respiratory activity using an MTT reduction assay. Tolfenpyrad caused a dose-dependent decrease in MTT reduction exclusively in *F. novicida* (Fig. 1E), indicating inhibition of electron transport-linked metabolism. Consistent with this interpretation, tolfenpyrad-resistant *F. novicida* isolates displayed attenuated MTT inhibition relative to wild type (Fig. S1). Collectively, these findings suggested that tolfenpyrad modulates ATP production through disruption of respiration, prompting us to investigate the *Francisella* electron transport chain (ETC) as a candidate target.

### Respiratory architecture underlying ATP production in F. novicida

To identify potential targets of inhibition, we mapped the architecture of the *F. novicida* electron transport chain (ETC) based on sequence homology, domain conservation, and functional annotation. Genomic analysis indicated the presence of a canonical 14-subunit NADH:ubiquinone oxidoreductase (complex I) containing conserved residues for flavin binding, iron-sulfur cluster coordination and proton translocation (Fig. 2)^26–28^. These features support its role as the primary entry point for NADH-derived electrons. *F. novicida* also encodes a non-proton pumping type II NADH dehydrogenase (NDH-2), which oxidizes NADH to transfer electrons into the ETC. Based on sequence conservation with NDH-2 homologs, this protein is unlikely to translocate protons, and therefore, does not directly contribute to the proton motive force generation^29^. Thus, complex I represents the major bioenergetic node for coupling NADH oxidation to energy conservation.

**Fig. 2.**
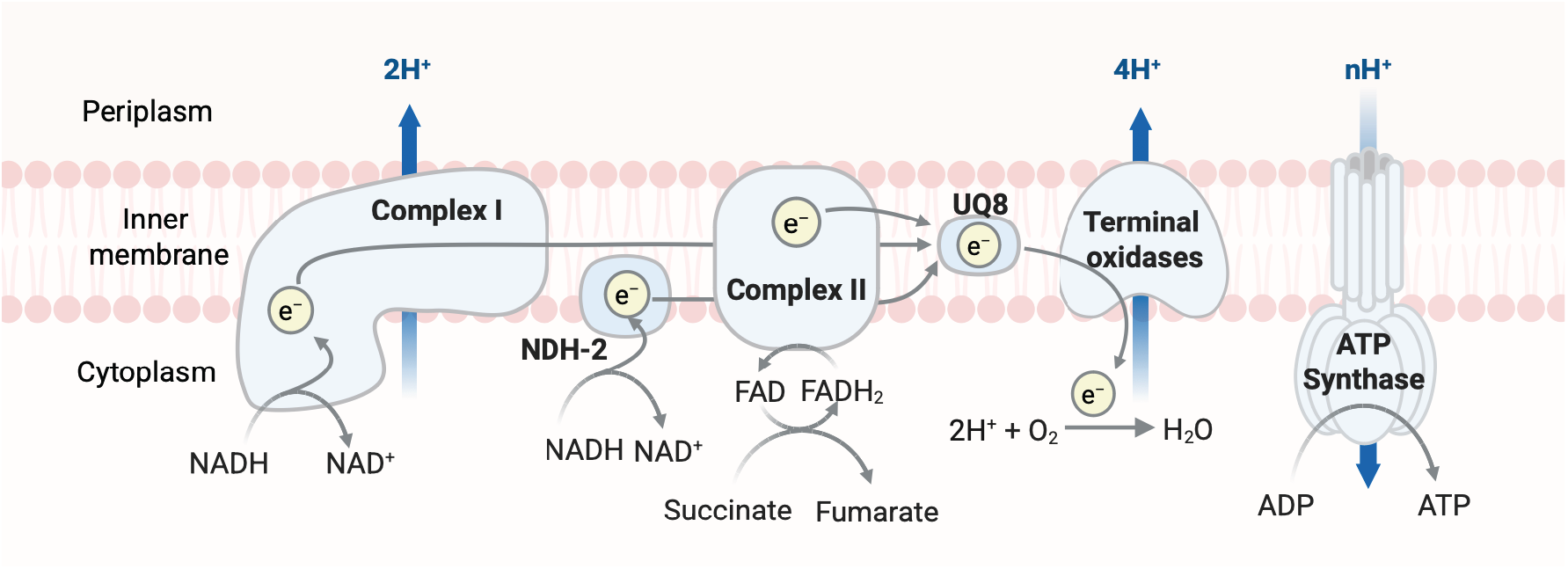
Schematic representation of the *F. novicida* electron transport chain. Predicted organization of the *F. novicida* electron transport chain based on functional annotation and sequence conservation. Electrons derived from NADH and succinate enter the quinone pool (UQ8) through Complex I, NDH-2, and Complex II and are transferred to terminal oxidases that reduce molecular oxygen. Proton translocation generates a proton motive force that drives ATP synthesis through the F_0_F_1_-ATP synthase.

*F. novicida* encodes succinate dehydrogenase (complex II), which oxidizes succinate and feeds electrons into the quinone pool without proton translocation. Previous studies indicate that ubiquinone-8 is the principal electron carrier in *F. novicida*, shuttling electrons from upstream dehydrogenases to terminal oxidases^30^. Similar to *E. coli*, no cytochrome bc_1_ complex (complex III) homologs were identified, indicating direct electron transfer from ubiquinone-8 to terminal oxidases^31^. Three terminal oxidases, including one cytochrome bo_3_-and two cytochrome bd-types, are predicted to reduce oxygen and contribute to proton motive force either by active proton pumping or by scalar proton consumption during oxygen reduction, thereby generating a transmembrane proton gradient. The resulting transmembrane proton gradient drives ATP synthesis via the F_0_F_1_ ATP synthase complex.

Together, this architecture defines a branched respiratory chain in which NADH- and succinate-derived electrons converge at the quinone pool before being transferred to terminal oxidases that further generate proton motive force to drive ATP synthesis. This organization provides a framework for identifying bioenergetic vulnerabilities and highlights complex I as a key point of susceptibility.

### Tolfenpyrad blocks oxygen consumption in F. novicida

*F. novicida* is an obligate aerobe that relies predominantly, if not exclusively, on oxygen as a terminal electron acceptor; thus, oxygen consumption provides a direct readout of ETC activity. We measured oxygen consumption to assess the impact of tolfenpyrad on respiratory electron transport. Untreated *F. novicida* exhibited robust oxygen consumption, which was abolished by tolfenpyrad (Fig. 3A). In contrast, this drug did not affect respiration in *E. coli* or *P. aeruginosa*, indicating species-specific inhibition. To exclude the possibility that reduced oxygen consumption reflected decreased bacterial viability, we enumerated CFU before and after respiration assays. CFU enumeration before and after respiration assays revealed no differences across conditions, confirming that altered oxygen consumption reflects metabolic inhibition rather than changes in bacterial number (Fig. 3B-3D). We next compared the sensitivity of these strains to rotenone, a canonical mammalian complex I inhibitor that does not inhibit the *E. coli* ETC due to structural divergence^32^. Rotenone selectively inhibited oxygen consumption in *F. novicida* but not in *E. coli* or *P. aeruginosa* (Fig. 3E). The concordance of *F. novicida* sensitivity to rotenone and tolfenpyrad suggested that drug susceptibility may correlate with complex I architecture. Together, these data indicate that tolfenpyrad acutely blocks respiratory activity in *F. novicida*, consistent with inhibition at the level of complex I.

**Fig. 3.**
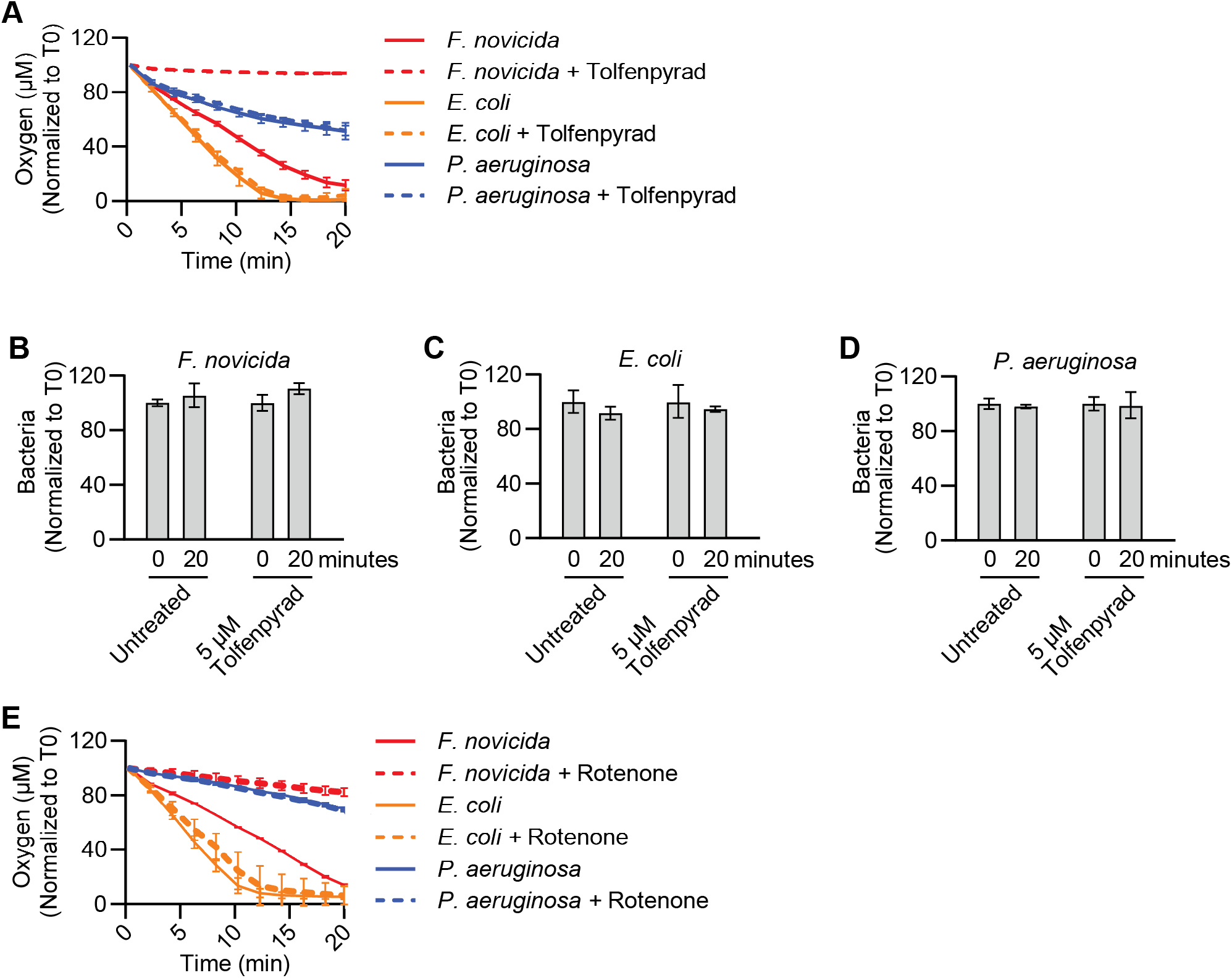
Tolfenpyrad selectively disrupts oxygen consumption in *F. novicida*. **(A)** Oxygen consumption in *F. novicida, E. coli*, and *P. aeruginosa* in the presence of 5 µM tolfenpyrad or vehicle control. Oxygen concentration was normalized to the beginning of the experiment (T0). **(B-D)** CFU enumeration of *F. novicida* **(B)**, *E. coli* **(C)**, and *P. aeruginosa* **(D)** before and after oxygen consumption measurements shown in (a). **(E)** Oxygen consumption following treatment with 5 µM rotenone or vehicle control. Data are presented as mean ± S.D. and are representative of three or more independent experiments performed with at least three technical replicates per condition.

### Respiratory inhibition by tolfenpyrad induces oxidative stress

Perturbation of electron transport can promote electron transfer to molecular oxygen, resulting in formation of reactive oxygen species (ROS). In particular, inhibition of NADH-dependent electron flow through complex I can increase electron leakage from respiratory complexes, generating superoxide that is subsequently converted to hydrogen peroxide^33^. Given the pronounced effects of tolfenpyrad on oxygen consumption in *F. novicida*, we asked whether respiratory inhibition by tolfenpyrad is accompanied by oxidative stress. Tolfenpyrad induced robust hydrogen peroxide production in *F. novicida*, comparable to the redox-cycling agent menadione (Fig. 4A)^34^. In contrast, tolfenpyrad did not induce ROS in *E. coli* or *P. aeruginosa*, although both species responded to menadione (Fig. 4B-C). Thus, the absence of ROS induction in these organisms is not due to an inability to generate or detect oxidative stress, but rather reflects their resistance to tolfenpyrad-mediated respiratory disruption. These findings demonstrate that tolfenpyrad-mediated respiratory blockade results in redox imbalance and oxidative stress specifically in *F. novicida*, providing an additional mechanistic consequence of electron transport chain inhibition.

**Fig. 4.**
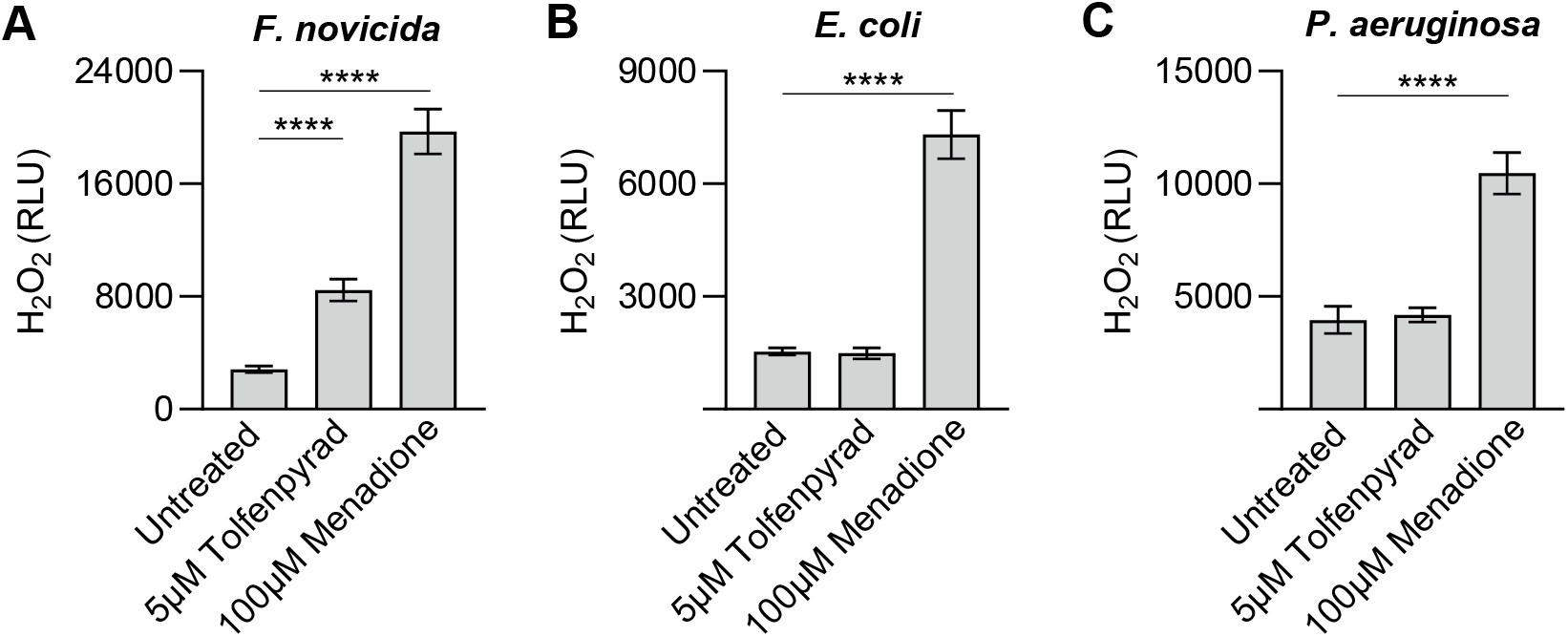
Tolfenpyrad induces reactive oxygen species in *F. novicida*. **(A-C)** Hydrogen peroxide production measured using ROS-Glo™ H_2_O_2_ assay (Promega) in *F. novicida* **(A)**, *E. coli* **(B)** and *P. aeruginosa* **(C)** after treatment with 5 µM tolfenpyrad, 100 µM menadione, or vehicle control. Data are presented as mean ± S.D. and are representative of three or more independent experiments performed with at least three technical replicates per condition. Statistical significance was determined using two-tailed unpaired t-tests. **** P ≤ 0.0001.

### Tolfenpyrad collapses proton motive force and exhibits pH-dependent activity

Because electron transport chain activity is coupled to proton translocation across the inner membrane, inhibition of respiratory electron flow would be expected to impair generation of proton motive force (PMF)^35^. To determine whether tolfenpyrad disrupts transmembrane proton gradients, we monitored PMF using the pH-sensitive fluorescent dye ACMA^36^. Untreated *F. novicida* rapidly quenched ACMA fluorescence, consistent with robust ETC-driven proton translocation (Fig. 5A). In contrast, increasing concentrations of tolfenpyrad progressively attenuated fluorescence quenching, indicating impaired proton gradient formation. At higher drug concentrations, fluorescence retention phenocopied treatment with SDS, which disrupts membrane integrity and abolishes proton gradients. These results demonstrate that tolfenpyrad disrupts ETC-dependent proton pumping, leading to collapse of the electrochemical gradient across the inner membrane. Notably, tolfenpyrad did not alter ACMA quenching in *E. coli* or *P. aeruginosa* (Fig. 5B, 5C), confirming that PMF disruption is selective for *F. novicida* and paralleling its species-specific effects on ATP production and oxygen consumption.

**Fig. 5.**
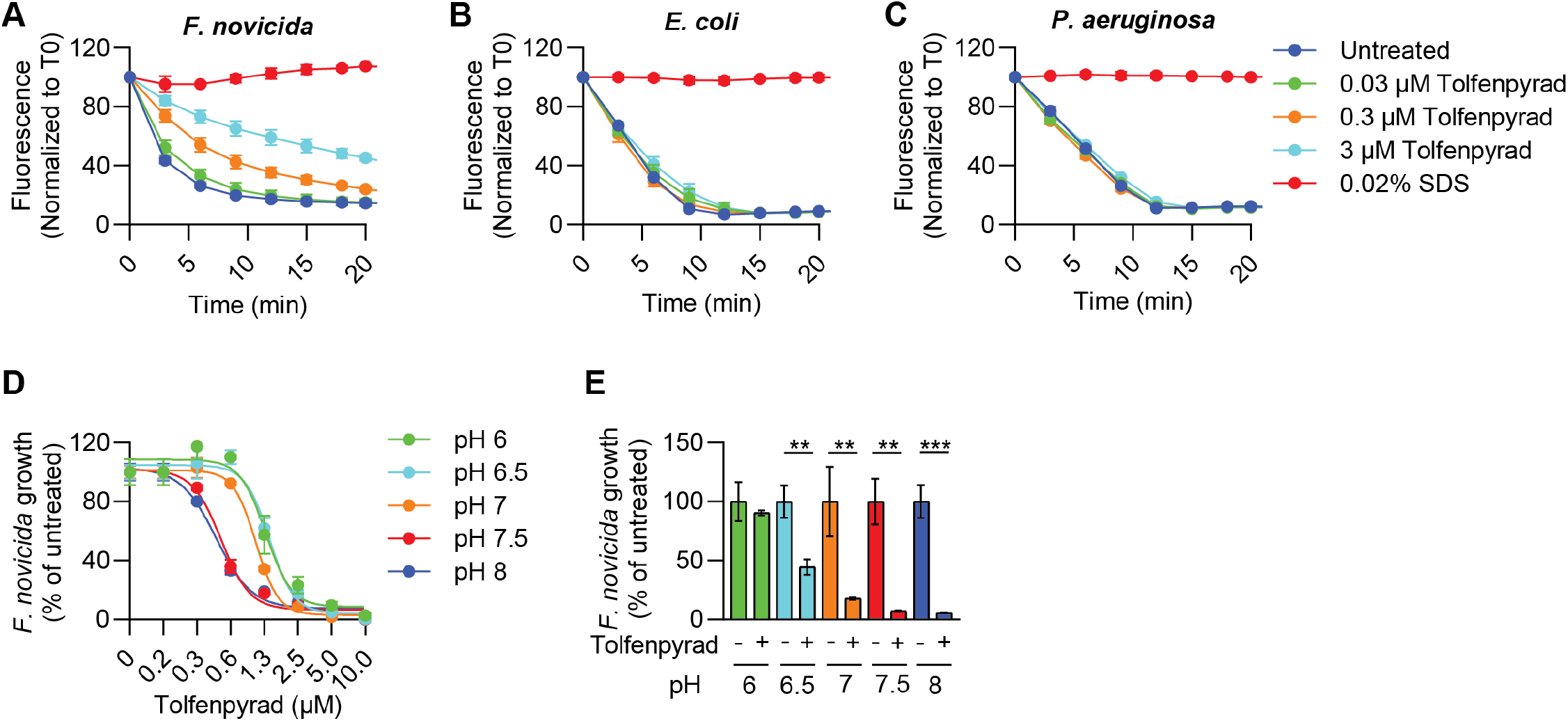
Tolfenpyrad disrupts proton gradient formation and pH-dependent activity in *F. novicida*. **(A-C)** Proton gradient measured by ACMA fluorescence quenching in membrane fractions isolated from *F. novicida* **(A)**, *E. coli* **(B)** and *P. aeruginosa* **(C)** following treatment with the indicated concentrations of tolfenpyrad. SDS (0.02%) was used as a membrane-disrupting control. Fluorescence values were normalized to T0. **(D)** Dose-response analysis of *F. novicida* growth measured at logarithmic phase by OD_600nm_ at the indicated extracellular pH values. **(E)** CFU quantification of *F. novicida* following 7 h treatment with 0.6 µM tolfenpyrad at the indicated pH values, normalized to untreated controls. Data are presented as mean ± S.D. and are representative of three or more independent experiments performed with at least three technical replicates per condition. Statistical significance was determined using two-tailed unpaired t-tests. ** P ≤ 0.01, *** P ≤ 0.001.

The PMF derives from a proton gradient across the inner membrane (∆pH), and environmental pH is predicted to influence bacterial sensitivity to respiratory inhibition^37^. We therefore tested whether extracellular pH modulates tolfenpyrad activity on bacterial cultures. Increasing the pH above neutral enhanced tolfenpyrad potency, lowering the IC_50_ from 1.5 μM to 0.5 μM, whereas lower pH reduced its effectiveness (Fig. 5D, 5E). Because elevated extracellular pH diminishes ΔpH, these findings indicate that partial PMF dissipation sensitizes *F. novicida* to respiratory inhibition. The enhanced activity of tolfenpyrad at elevated pH therefore suggests that partial dissipation of proton motive force sensitizes *F. novicida* to further disruption of respiratory energy transduction. Together with the ACMA measurements, these findings reinforce the conclusion that tolfenpyrad impairs ETC-dependent proton pumping and compromises bioenergetic homeostasis in *F. novicida*.

### Tolfenpyrad selectively inhibits complex I-dependent respiration

Complex I catalyzes the oxidation of NADH and transfers electrons to ubiquinone, generating NAD^+^ and initiating electron flux through the respiratory chain. Because inhibition of this step perturbs intracellular redox balance, we quantified NAD:NADH ratios in bacteria treated with tolfenpyrad. Untreated *F. novicida* maintained an NAD^+^:NADH ratio of ∼4, indicative of active respiratory metabolism (Fig. 6A). Even at the IC_10_ (0.3 µM), tolfenpyrad decreased this ratio by >60%, with a greater effect at higher concentrations, reflecting a shift toward a more reduced state and impaired NADH oxidation. Potassium cyanide produced a similar response in *F. novicida*, supporting disruption of intracellular redox homeostasis through inhibition of respiratory electron flow. In contrast, the NAD⁺:NADH ratio in *E. coli* was unaffected by tolfenpyrad but decreased in response to ZnCl_2_, consistent with inhibition of complex I activity (Fig. 6B)^38^. These findings demonstrate that tolfenpyrad rapidly disrupts intracellular redox homeostasis in a manner consistent with inhibition of respiratory electron flow.

**Fig. 6.**
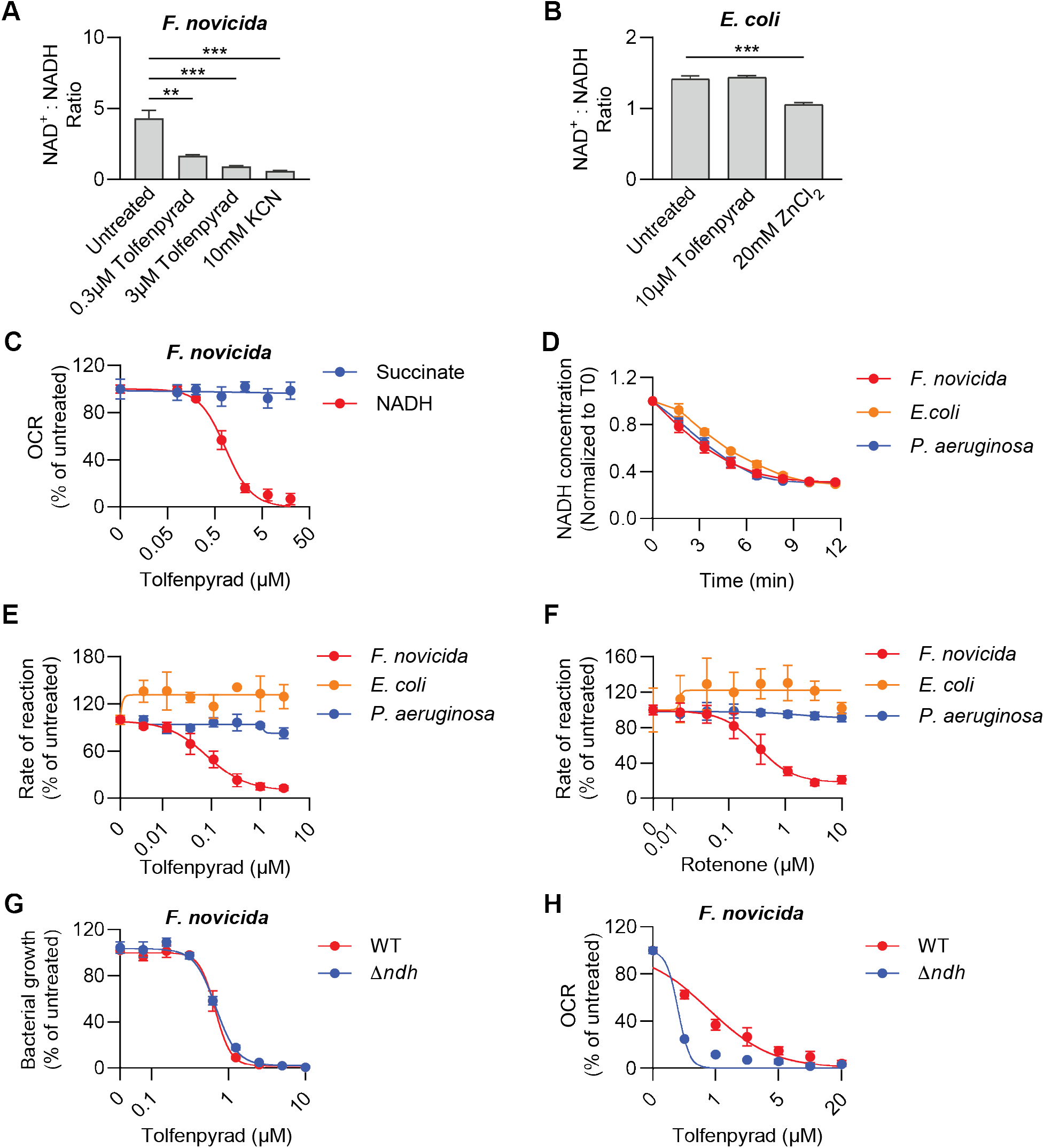
Tolfenpyrad inhibits Complex I-dependent NADH oxidation in *F. novicida*. **(A, B)** NAD^+^:NADH ratios in *F. novicida* **(A)** and *E. coli* **(B)** following treatment with the indicated compounds for 1 h. **(C)** Oxygen consumption rates (OCR) of *F. novicida* membrane fractions supplied with 2.5 mM NADH or 30 mM succinate in the presence of the indicated concentrations of tolfenpyrad. **(D)** NADH oxidation measured by loss of absorbance at 340 nm in membrane fractions isolated from *F. novicida, E. coli*, and *P. aeruginosa*. **(E, F)** Quantification of NADH oxidation rates following treatment with the indicated concentrations of tolfenpyrad **(E)** or rotenone **(F). (G)** Dose-response analysis of wild-type *F. novicida* (WT) and the Δ*ndh* mutant based on OD_600nm_ measurements. **(H)** OCR measurements in membrane fractions harvested from wild-type *F. novicida* (WT) and the Δ*ndh* mutant supplied with 2.5 mM NADH as the substrate after a 10 min exposure with the indicated concentrations of tolfenpyrad. Data are presented as mean ± S.D. and are representative of three or more independent experiments performed with at least three technical replicates per condition. ** P ≤ 0.01.

To determine whether tolfenpyrad impairs NADH-dependent electron entry into the electron transport chain or affects downstream steps, we performed substrate-specific respiration assays using liposomes derived from *F. novicida, E. coli*, and *P. aeruginosa*. When *F. novicida* membranes were supplied with NADH as an electron donor, tolfenpyrad produced dose-dependent inhibition of oxygen consumption, consistent with suppression of NADH-linked respiratory activity (Fig. 6C). In contrast, oxygen consumption driven by the substrate for complex II, succinate, was unaffected by tolfenpyrad, excluding complex II as a target and suggesting that tolfenpyrad selectively interferes with NADH-dependent electron entry into the respiratory chain.

We next measured NADH oxidation *in vitro* in an enzymatic assay. NADH absorbs light at 340 nm, whereas NAD⁺ does not, allowing kinetic monitoring of NADH consumption by measuring a decrease in absorbance. Membrane fractions isolated from all three species exhibited comparable baseline NADH oxidation rates in the absence of inhibitor, indicating similar catalytic capacity under the assay conditions (Fig. 6D). Tolfenpyrad selectively inhibited NADH oxidation in *F. novicida* membranes but not in *E. coli* or *P. aeruginosa* (Fig. 6E). To determine whether this inhibition phenocopies canonical complex I blockade, we compared tolfenpyrad with rotenone, a well-characterized complex I inhibitor. Rotenone strongly inhibited NADH oxidation in *F. novicida* membranes but had no detectable effect on membranes derived from *E. coli* or *P. aeruginosa* (Fig. 6F). Thus, like Rotenone, tolfenpyrad inhibits NADH-dependent respiratory activity in a species-selective manner, mirroring its effects on oxygen consumption in intact cells. In addition to complex I, the *F. novicida* ETC contains a second NADH-oxidizing enzyme, NDH-2, which also transfers electrons to ubiquinone. To distinguish between NADH oxidation by complex I or NDH-2, we tested the tolfenpyrad sensitivity of a Δ*ndh* mutant to tolfenpyrad. Drug sensitivity of the Δ*ndh* mutant was identical to wild type, excluding it as the primary target (Fig. 6G). Consistent with NDH-2 being dispensable for sensitivity to tolfenpyrad, NADH-dependent respiration by membranes derived from the Δ*ndh* mutant was more sensitive to tolfenpyrad than wild type (Fig. 6H). This enhanced inhibition is consistent with complex I serving as the predominant remaining NADH-oxidizing enzyme in the mutant background, thereby increasing reliance on the tolfenpyrad-sensitive pathway.

A common mechanism by which respiratory inhibitors impair complex I activity is by preventing electron transfer from NADH to ubiquinone through direct competition at the ubiquinone-binding pocket. If tolfenpyrad inhibits complex I by occupying or sterically blocking this site, increasing the availability of ubiquinone would be expected to competitively overcome inhibition and partially restore respiratory activity. To test this possibility, we supplemented *F. novicida* cultures with exogenous ubiquinone prior to drug exposure and assessed sensitivity to tolfenpyrad. Pre-incubation with excess ubiquinone did not alter bacterial susceptibility to tolfenpyrad, indicating that increased substrate availability does not rescue inhibition (Fig. S2A).

Similarly, when bacteria were first exposed to tolfenpyrad and subsequently supplied with excess ubiquinone, no reduction in drug sensitivity was observed (Fig. S2B). Thus, elevating ubiquinone concentrations either before or after inhibitor exposure failed to modulate the antibacterial effects of tolfenpyrad. Together, these findings argue against a competitive interaction between tolfenpyrad and the ubiquinone-binding site of complex I. Instead, the data suggest that tolfenpyrad inhibits NADH-dependent respiration through a mechanism distinct from blockade of ubiquinone binding or reduction, consistent with inhibition occurring at an alternative functional or structural region of the enzyme complex.

### Structural modeling prioritizes NuoM subunit as a candidate interaction site

To identify potential tolfenpyrad interaction sites, we performed unbiased blind docking against AlphaFold-predicted structures of individual subunits of the *F. novicida* complex I. No binding pockets were predefined, enabling an agnostic assessment of potential ligand-binding regions across the full protein surfaces. Several complex I subunits, including NuoF, NuoH, NuoL, and NuoM produced favorable docking scores (ΔG < -8.0 kcal/mol). We then repeated the analysis with NK-06, a tolfenpyrad analogue that we previously demonstrated has enhanced antibacterial activity^23^. Among the candidate subunits, NuoM showed the largest increase in predicted binding affinity with NK-06 (ΔG = -9.1 kcal/mol, Table 1), consistent with our previous finding that nuoM is mutated in multiple tolfenpyrad-resistant isolates^39^.

**Table 1.** Blind docking scores of tolfenpyrad and NK-06 across *F. novicida* Complex I subunits. Values represent AutoDock Vina docking scores (kcal/mol).

| Complex I Subunit | Tolfenpyrad | NK-06 |
| --- | --- | --- |
| NuoA | -6.53 | -6.53 |
| NuoB | -7.13 | -6.44 |
| NuoC | -7.18 | -7.33 |
| NuoD | -7.06 | -7.25 |
| NuoE | -6.55 | -7.08 |
| NuoF | -8.68 | -8.43 |
| NuoG | -7.51 | -7.61 |
| NuoH | -8.72 | -8.17 |
| NuoI | -6.70 | -7.31 |
| NuoJ | -6.64 | -6.76 |
| NuoK | -7.15 | -6.81 |
| NuoL | -8.36 | -8.97 |
| NuoM | -7.84 | -9.10 |
| NuoN | -7.10 | -7.36 |

Docking predicted that tolfenpyrad was able to bind a hydrophobic tunnel within *F. novicida* NuoM, positioning its aromatic moieties along a membrane-embedded groove within the subunit while orienting the pyrazole group towards the surface (Fig. 7A). NK-06 occupied the same region within NuoM, further supporting this site as a candidate interaction region. Comparison of the *F. novicida* NuoM model to the solved structure for *E. coli* NuoM revealed strong overall similarity between these proteins (RMSD for Cα < 1.0 Å) (Fig. 7B). Although these orthologs share a high degree of structural homology, the tunnel identified in *F. novicida* NuoM was absent in the corresponding region of *E. coli*, correlating with the lack of experimentally observed tolfenpyrad inhibition in *E. coli* (Fig. 7C). These differences provide a potential structural explanation for species-specific inhibition and identify NuoM as a candidate interaction site.

**Fig. 7.**
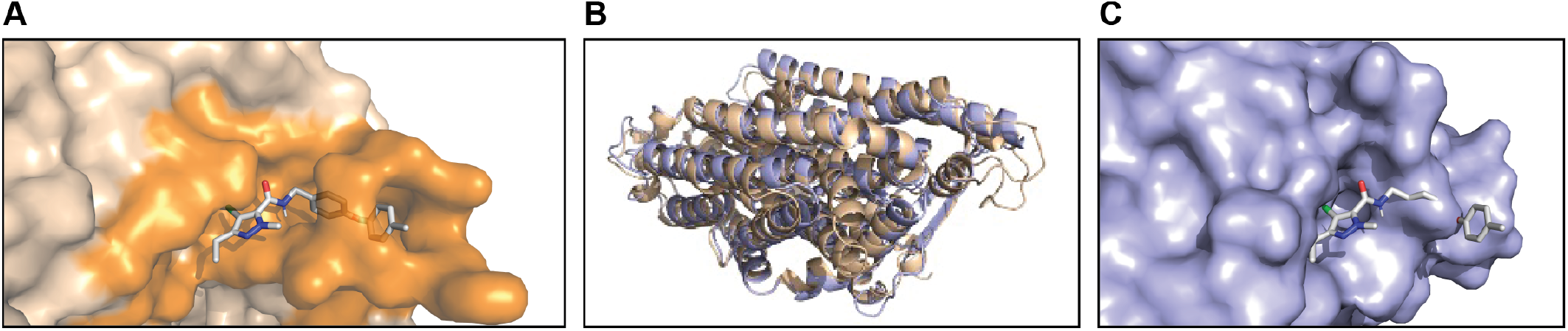
Structural modeling identifies NuoM as a candidate tolfenpyrad interaction site. **(A)** Blind docking localized tolfenpyrad to a tunnel region within *F. novicida* NuoM. Tolfenpyrad is shown as sticks and the tunnel region is shown as a semi-transparent surface. **(B)** Structural alignment of an AlphaFold-predicted NuoM model from *F. novicida* (wheat) with the solved structure from *E. coli* (light blue, PDB: 3M9C) demonstrates a high degree of overall structural similarity between the two orthologs. **(C)** Surface representation of *E. coli* NuoM with the tolfenpyrad docking pose from the aligned *F. novicida* structure superimposed, illustrating the absence of the tunnel architecture observed in *F. novicida*. Structural models were visualized using PyMOL.

Collectively, these results demonstrate that tolfenpyrad selectively targets *Francisella* complex I, collapsing electron transport, proton motive force, and ATP production while inducing oxidative stress. Structural divergence in the NuoM subunit provides a mechanistic basis for this specificity and reveals bacterial complex I as a previously unrecognized therapeutic vulnerability in *Francisella* metabolism.

## DISCUSSION

ATP generation through respiration is a cornerstone of bacterial survival, yet the electron transport chains of many intracellular pathogens remain poorly characterized^40^. Here, we identify respiratory complex I as a selective and druggable vulnerability in *Francisella* and establish the small molecule tolfenpyrad as a species-specific inhibitor of NADH-dependent electron transport. We show that tolfenpyrad disrupts electron flow, collapses proton motive force, depletes intracellular ATP, and induces ROS production, collectively driving bacteriostasis in *F. novicida*. These findings demonstrate that disruption of respiratory energy transduction is sufficient to impair *Francisella* viability and positions complex I as a previously unrecognized antimicrobial target in this genus.

Multiple lines of evidence support a model in which tolfenpyrad inhibits NADH-dependent entry into the electron transport chain at the level of complex I^41,42^. Tolfenpyrad selectively blocks NADH-driven respiration without affecting succinate-dependent electron flow, rapidly perturbs intracellular NAD⁺:NADH ratios, and phenocopies the effects of rotenone^43^. Genetic and biochemical analyses further exclude NDH-2 as the primary target, while increased sensitivity in the Δ*ndh* mutant is consistent with enhanced reliance on complex I activity. Together, these data demonstrate that tolfenpyrad inhibits complex I-mediated electron transfer rather than downstream respiratory components or alternative NADH dehydrogenases.

Despite broadly conserved respiratory architectures among γ -proteobacteria, this functional response was specific to *Francisella*. Neither *E. coli* nor *P. aeruginosa* exhibited detectable sensitivity to tolfenpyrad across assays of respiration, ATP production, or ROS induction^22,43^. This selectivity extends beyond tolfenpyrad, as *Francisella* displays heightened sensitivity to multiple electron transport chain inhibitors, including rotenone, clofazimine, and benzarone, which have limited activity against other Gram-negative bacteria^43^. Notably, *Francisella* rely strictly on oxygen as a terminal electron acceptor and lack alternative respiratory pathways, potentially increasing its dependence on complex I-driven proton translocation and rendering it uniquely susceptible to inhibition^30,31^.

Our structural modeling provides a mechanistic basis for this selectivity. Docking simulations predict that tolfenpyrad binds within a tunnel in the membrane subunit NuoM of *F. novicida* complex I, whereas the corresponding region in *E. coli* lacks this tunnel. Tolfenpryad’s insecticidal activity is also associated with inhibiting complex I, highlighting a conserved mechanistic vulnerability while underscoring structural differences unique to *Francisella*. These findings are consistent with prior identification of resistance-inducing mutations in *nuoM* and suggest that structural divergence within this subunit governs species-specific susceptibility^22^. NuoM is part of a conserved set of antiporter-like subunits that couple redox reactions in the peripheral arm of complex I to proton translocation across the membrane^44,45^. Binding of tolfenpyrad in this region may disrupt conformational coupling required for proton pumping, thereby uncoupling electron transfer from energy conservation. Structural studies of mitochondrial complex I have identified three rotenone binding sites, including one in ND4, the homolog of bacterial NuoM. Consistent with this, the sensitivity of *F. novicida* to rotenone supports the conclusion that this membrane-arm region may represent a potential inhibitor-sensitive site in *Francisella*^46^.

Disruption of complex I activity had profound consequences for *Francisella* physiology. Tolfenpyrad-induced inhibition of electron transport resulted in rapid ATP depletion and collapse of proton motive force, consistent with the central role of complex I in maintaining bioenergetic homeostasis^45^. In parallel, inhibition of NADH oxidation shifted intracellular redox balance toward a more reduced state and triggered robust ROS production. This combination of energetic failure and oxidative stress likely underlies the observed bacteriostatic phenotype. The strong pH dependence of tolfenpyrad activity further supports a bioenergetic mechanism, as increasing extracellular pH reduces the ∆pH component of the proton motive force and sensitizes bacteria to further disruption^37^. These findings highlight the importance of environmental context in modulating the efficacy of respiratory inhibitors and suggest that host niche conditions may influence antimicrobial activity *in vivo*^47^.

Despite the pronounced effects of tolfenpyrad on respiratory activity and bioenergetic homeostasis, the compound exhibited primarily bacteriostatic rather than bactericidal activity under the conditions tested. One possible explanation is that residual respiratory activity or alternative metabolic pathways may partially sustain viability despite impaired complex I-dependent electron transport. In addition, bacteria experiencing energetic stress can adopt low-metabolic or growth-arrested states that enhance survival during respiratory inhibition^48^. The ability of *F. novicida* to tolerate substantial disruption of ATP production and proton motive force may therefore reflect physiological adaptation to metabolic stress rather than complete bioenergetic collapse. Future studies will be required to determine whether prolonged exposure, combinatorial therapies, or host-associated stresses convert this phenotype into bactericidal activity.

The selectivity of tolfenpyrad is particularly notable given the functional conservation of complex I across bacteria and mitochondria^49,50^. While complex I is considered a challenging antimicrobial target, our findings demonstrate that species-specific structural features can be exploited to achieve selective inhibition^41,51^. The identification of NuoM as a candidate interaction site is especially intriguing, as it implicates a membrane-embedded, proton-translocating subunit rather than the canonical redox-active regions typically targeted by inhibitors^44,45,52^. Targeting such noncanonical functional domains may provide a strategy to achieve both potency and specificity while minimizing off-target effects^53^.

From a translational perspective, these findings position respiratory metabolism as a promising avenue for therapeutic development against *Francisella*. The ability of tolfenpyrad to selectively impair bacterial energetics and induce oxidative stress suggests potential utility as a direct antimicrobial or as part of combination therapies. Disruption of proton motive force is known to influence antibiotic uptake and efficacy, raising the possibility of synergistic interactions with existing treatments^43^. Although tolfenpyrad itself has known toxicity associated with complex I inhibition in eukaryotic systems, we have identified derivatives that retain antibacterial activity while exhibiting reduced toxicity in macrophages, providing a foundation for further optimization^23,54^. Future studies will be required to evaluate the *in vivo* efficacy of these compounds and to determine whether similar vulnerabilities can be exploited in other intracellular pathogens with comparable respiratory architectures.

Several limitations of this study should be noted. Although our genetic, biochemical, and physiological data strongly support complex I as the primary target of tolfenpyrad in *Francisella*, direct structural or biophysical evidence of inhibitor binding remains to be established. In particular, the proposed interaction with the membrane subunit NuoM is based primarily on computational modeling together with indirect genetic evidence and will require future validation through structural and mechanistic studies. While previous work demonstrated that tolfenpyrad exhibits selective antibacterial activity against *Francisella* and retains activity in macrophage infection models, the broader spectrum of susceptibility across related bacterial species and the *in vivo* therapeutic potential of respiratory inhibition remain incompletely defined. Future studies examining resistance mechanisms, target engagement, and efficacy in animal models will further clarify the utility of targeting respiratory metabolism in Francisella.

In summary, we demonstrate that selective inhibition of complex I disrupts *Francisella* bioenergetics and reveals a previously unappreciated vulnerability in its respiratory machinery. These findings expand the repertoire of targetable pathways in bacterial pathogens and establish a framework for exploiting species-specific features of central metabolism for antimicrobial discovery.

## METHODS

### Bacterial strains and culture conditions

*F. novicida* Utah 112 (U112, NR-13) was obtained from the Biodefense and Emerging Infections Research Resources Repository (BEI resources) at the National Institute of Allergy and Infectious Diseases (NIAID), an institute of the National Institutes of Health (NIH). *P. aeruginosa* PAO1 and *E. coli* P4 were obtained from Joseph Mougous (Yale University) and Ricardo C. Chebel (University of Florida), respectively. *F. novicida* was maintained on tryptic soy agar or broth containing 0.1% cysteine (TSAC/TSBC). *E. coli* and PAO1 were cultured in the same media lacking cysteine (TSA/TSB).

An in-frame, markerless ∆*ndh* (FTN_0912) deletion mutant of *F. novicida* was generated by amplifying ∼500–600 bp regions upstream and downstream of *ndh* using the following primers: upstream forward (cggccagtgccaagctgaagctacagaagaggagattatcg) and reverse (cttaattttcaaattttattatttttctcatactttccccttac), and downstream forward (gaaaaataataaaatttgaaaattaagaggtctattcac) and reverse (caggaaacagctatgacatgattaggtaatagacctgatgaaa acttcc). The fragments were spliced by overlap extension PCR, cloned into the pEX18km vector by Gibson assembly, and introduced into *F. novicida* for allelic replacement, as described previously^55^. The resulting mutant retained a 30 bp remnant of *ndh*, as confirmed by colony PCR and Sanger sequencing.

### Bacterial growth quantification

To monitor the bacterial growth kinetics, bacteria were grown overnight on TSAC or TSA agar plates. Broth cultures were inoculated from freshly streaked agar plates into TSBC or TSB, adjusted to an optical density at 600 nm (OD_600nm_) of 0.001, sealed with breathable membranes (Breathe-Easy) and incubated at 37°C while shaking. Samples were collected at 60-min intervals, serially diluted in TSBC, and plated for CFU enumeration using standard methods^56^. After 3 h, tolfenpyrad was added at the indicated concentration, and samples were subsequently collected every 30 min for CFU enumeration.

Bacterial growth in the indicated media, including different pH conditions and TSBC, was monitored using a plate reader. Overnight cultures were diluted 1:100 into the appropriate medium and normalized to an OD600 of 0.05. Bacteria were dispensed into 96-well plates (n = 4) and treated with a titration of tolfenpyrad. Plates were sealed with a breathable membrane, and growth was monitored by measuring absorbance at 600 nm every 20 min using a BioTek Synergy HTX multimode plate reader. Dose-response curves were analyzed and plotted at the time point when bacteria reached to logarithmic phase. In parallel, bacterial growth across five pH conditions in TSBC was independently quantified by CFU enumeration following 7 h incubation with 0.6 µM tolfenpyrad or 0.04% vehicle control.

### Measurement of Respiratory Activity

Log-phase bacteria were normalized to OD_600nm_ = 0.25 (*F. novicida, E. coli*) or 0.5 (*P. aeruginosa*) and added to 96-well plates containing tolfenpyrad at the indicated concentrations. After 15 min at 37°C with shaking, MTT was added to 0.6 mg/mL and incubated for 30 min. DMSO was then added (1:1) to solubilize formazan, and respiratory activity was measured at A_540nm_ using a BioTek Synergy HTX plate reader.

### ATP quantification

ATP levels were measured using the ATP-Glo™ Assay Kit (Biotium) according to the manufacturer’s instructions. Exponential-phase bacteria were adjusted to OD_600nm_ = 0.5, treated with tolfenpyrad in 96-well plates, and sampled immediately before treatment and after 45 min at 37°C with shaking. Samples from the same wells were simultaneously collected for CFU enumeration. Samples were mixed with reagent in opaque 384-well plates, and luminescence (RLU) was measured using a plate reader. ATP levels were normalized to CFU and expressed relative to untreated controls (set to 100%).

### Quantification of Reactive oxygen species production

Reactive oxygen species (ROS) was quantified using the ROS-Glo^™^ H_2_O_2_ Assay (Promega) according to the manufacturer’s instructions. Exponential-phase bacteria were normalized to OD_600nm_ = 0.05 in TSBC/TSB, plated in white-sided, clear-bottom 384-well plates, and treated with 5 µM tolfenpyrad, 100 µM menadione, or 0.04% DMSO. H_2_O_2_ substrate was added to 25 µM, and plates were incubated at room temperature for 3 h. Detection solution was then added (1:1), and luminescence was measured using a microplate reader.

### Measurement of NAD^+^: NADH ratio in bacterial cultures

Exponential-phase bacteria were normalized to OD_600nm_ = 0.1 (*F. novicida*) or 0.25 (*E. coli*), plated in 96-well plates, and treated with the indicated compound or 0.04% DMSO. After 1 h at 37°C with shaking, NAD⁺:NADH ratios were measured using the NAD/NADH-Glo™ Assay Kit (Promega) according to the manufacturer’s instructions.

### Bacterial membrane isolation

Bacterial membrane fractions were prepared as described previously^57^. Briefly, cells were grown from overnight cultures in 1 L of TSB or TSBC to OD_600nm_ = 1.5 and harvested by centrifugation (6,000 × G, 15 min). Pellets were resuspended to 10% (w/v) in buffer (10 mM Tris-HCl pH 7.0, 1 mM EDTA, 1 mM DTT, 1 mM PMSF, 15% glycerol) and treated with lysozyme (0.5 mg/mL) on ice for 30 min. Cells were disrupted by sonication (∼15 cycles of 20 s; 1 s on/1 s off) and cleared by centrifugation (23,400 × g, 15 min, 4°C). Membranes were pelleted by ultracentrifugation (256,600 × g, 30 min), resuspended in buffer, and protein concentration was determined using the Pierce™ Bradford assay. Samples were stored at ™80°C until use.

### Quantification of oxygen consumption

Oxygen consumption was measured under conditions non-permissive for growth. Bacterial cultures were washed, adjusted to OD_600nm_ = 0.2, treated with tolfenpyrad (5 µM), rotenone (5 µM), or DMSO (0.04%) and 100 µL was added to 96-well plates. Oxygen levels were recorded for 30 min at 24 ± 1 °C using the Resipher (Lucid Scientific). CFU counts were measured from matched samples before and after the assay.

Oxygen consumption rates (OCR) were measured in isolated membranes from wild-type *F. novicida* and Δ*ndh*. Membranes were diluted in assay buffer (10 mM potassium phosphate, 1 mM EDTA, 5 mM MgCl_2_), plated in 96-well plates, and treated with the indicated concentrations of tolfenpyrad. Reactions were initiated with 2.5 mM NADH or 30 mM succinate, and oxygen consumption was continuously monitored using the Resipher. OCR was calculated as the change in oxygen concentration over time and normalized to membrane protein concentration.

### Quantification of proton gradient in bacterial membranes

Proton gradients were measured using ACMA fluorescence. Membrane fractions were diluted in buffer (50 mM MOPS pH 7.3, 50 mM KCl, and 10 mM MgCl_2_), and added to 96-well plates at 80 µg/mL (*F. novicida* and *E. coli*) or 160 µg/mL (*P. aeruginosa*). Samples were with the indicated compounds, supplemented with 2 µM ACMA, and initiated by addition of 150 µM NADH. Fluorescence (Ex/Em 360/460) was monitored over time using a microplate reader as a measure of proton motive force.

### NADH oxidation by bacterial membranes

To measure membrane-associated complex I activity, bacterial membranes were diluted with enzymatic assay buffer (10 mM potassium phosphate, 1 mM EDTA, 5 mM MgCl_2_) in a 96-well plate at final concentrations of 25 µg/mL for *F. novicida*, 62 µg/mL for *E. coli*, and 100 µg/mL for *P. aeruginosa*. Concentrations were optimized to yield similar rates of NADH oxidation. Wells were treated with the indicated concentrations of compound or 0.04% DMSO, and reactions were initiated with 150 µM NADH. NADH oxidation was monitored by the decrease in A_340nm_ in a plate reader.

### Molecular docking studies

Structural models of the 14 *F. novicida* strain U112 complex I subunits, NuoA - NuoN, were obtained from the AlphaFold Protein Structure Database. The corresponding AlphaFold entries were A0Q8H2, A0Q8H1, A0Q8H0, A0Q8G9, A0Q8G8, A0Q8G7, A0Q8G6, A0Q8G5, A0Q8G4, A0Q8G3, A0Q8G2, A0Q8G1, A0Q8G0, and A0Q8F9. Receptor models and tolfenpyrad were prepared for docking using AutoDock Tools (version 1.5.6) and converted to PDBQT format. Blind docking was performed using AutoDock Vina (version 1.2.3), with docking grids centered on each receptor and sized to encompass the full predicted structure^58,59^. Receptors were treated as rigid, whereas ligand torsions were allowed to rotate. Docking outputs were ranked according to Vina score.

### Statistical analysis and rigor of the data

All experiments were performed with at least three technical and three biological replicates. Statistical analysis and graphing were performed using GraphPad Prism (Version 10.4.1). Data are presented as means +/-standard deviation or standard error, as indicated and appropriate. Statistical significance was determined by using two-tailed unpaired t-tests and p-values are indicated in the figure legends.

## ACKNOWLEDGEMENTS

This study was supported by grants from the Higher Education Commission of Pakistan (F2022-442 to N.K.) and the United States National Institutes of Health (1R21AI204394 to A.E.), University of Florida Office of Research and College of Veterinary Medicine.

## SUPPLEMENTARY FIGURES

**Fig. S1.**
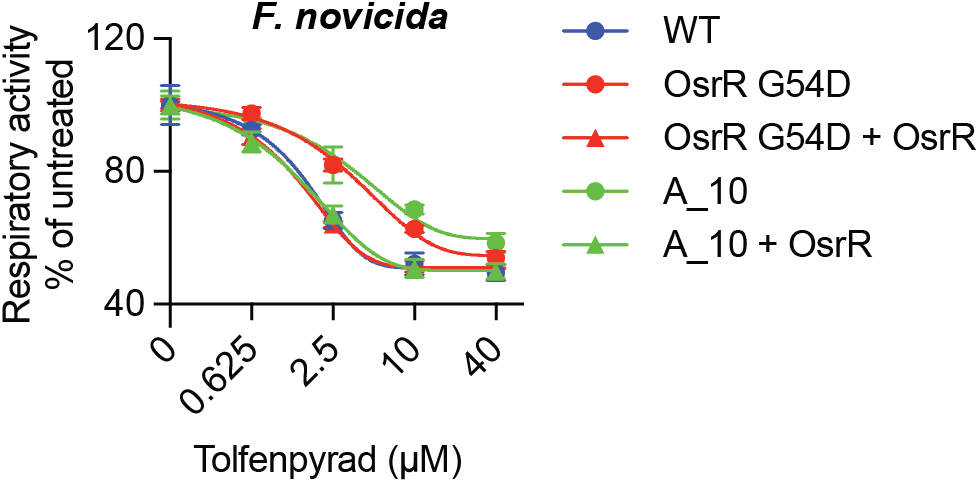
Tolfenpyrad-resistant *F. novicida* strains exhibit reduced respiratory inhibition. Respiratory activity measured by MTT reduction in wild-type (WT) *F. novicida*, and a tolfenpyrad-resistant mutant and isolate (OsrR G54D and A_10, respectively), as well as the mutants expressing wild-type OsrR following treatment with the indicated concentrations of tolfenpyrad. Data are presented as mean ± S.D. and are representative of three or more independent experiments performed with at least three technical replicates per condition.

**Fig. S2.**
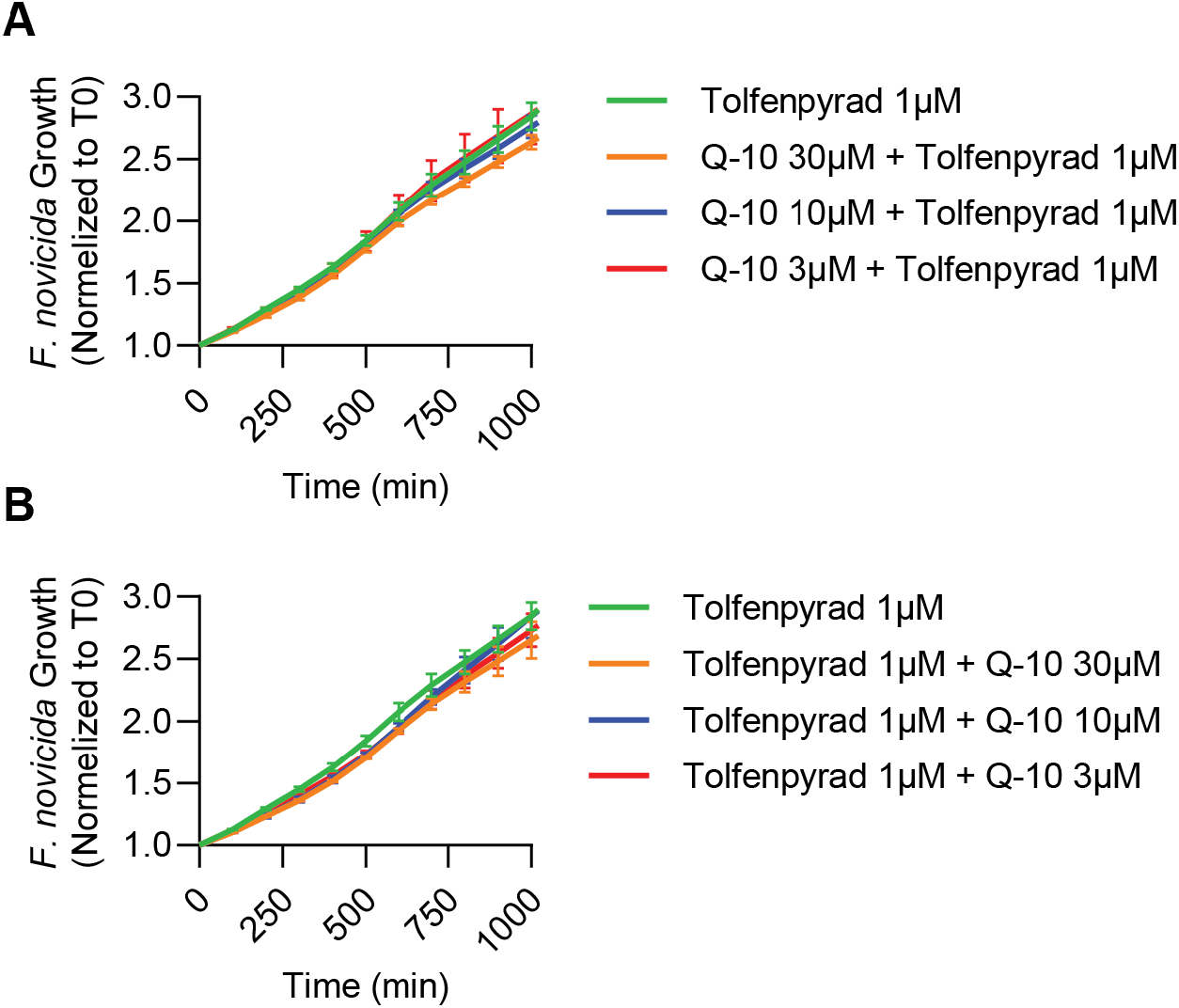
Exogenous ubiquinone does not rescue tolfenpyrad-mediated growth inhibition. **(A)** Growth of *F. novicida* following pre-incubation with the indicated concentrations of coenzyme Q10 prior to treatment with 1 µM tolfenpyrad. **(B)** Growth of *F. novicida* following treatment with 1 µM tolfenpyrad before addition of coenzyme Q10.

## Notes

### Competing Interest Statement

The authors have declared no competing interest.

